# SWS2 visual pigment evolution as a test of historically contingent patterns of plumage color evolution in Warblers

**DOI:** 10.1101/013573

**Authors:** Natasha I. Bloch, James M. Morrow, Belinda S.W. Chang, Trevor D. Price

## Abstract

Distantly related clades that occupy similar environments may differ due to the lasting imprint of their ancestors – historical contingency. The New World warblers (Parulidae) and Old World warblers (Phylloscopidae) are ecologically similar clades that differ strikingly in plumage coloration. We studied genetic and functional evolution of the short-wavelength sensitive visual pigments (SWS2 and SWS1) to ask if altered color perception could contribute to the plumage color differences between clades. We show SWS2 is short-wavelength shifted in birds that occupy open environments, such as finches, compared to those in closed environments, including warblers. Sequencing of opsin genes and phylogenetic reconstructions indicate New World warblers were derived from a finch-like form that colonized from the Old World 15-20Ma. During this process the SWS2 gene accumulated 6 substitutions in branches leading to New World warblers, inviting the hypothesis that passage through a finch-like ancestor resulted in SWS2 evolution. In fact, we show spectral tuning remained similar across warblers as well as the finch ancestor. Results reject the hypothesis of historical contingency based on opsin spectral tuning, but point to evolution of other aspects of visual pigment function. Using the approach outlined here, historical contingency becomes a generally testable theory in systems where genotype and phenotype can be connected.

## Introduction

Historical contingency refers to the lasting impression an ancestral form leaves on its descendants (Gould 2002). Even in the face of identical selection pressures, differences in ancestors will generally drive evolution along different trajectories (Gould 2002; Losos and Ricklefs 2009; Prunier et al. 2012). A role for contingency is most easily assessed in comparisons among species occupying similar environments (Losos and Ricklefs 2009). Such species are often convergent in many features, including spectacular examples of morphological convergence that are present between distantly related species (Fain and Houde 2004; Alvarado-Cárdenas and Martínez-Meyer 2013), as well as closer relatives (Losos and Ricklefs 2009; Mahler et al. 2013). However, convergence is rarely complete. If residual differences between environments can be ruled out as the cause (Alvarado-Cárdenas and Martínez-Meyer 2013), the failure to converge should reflect effects of the genetic and phenotypic make up of ancestors on the subsequent radiation, i.e., historical contingency (Schluter 1986; Losos and Ricklefs 2009; Prunier et al. 2012).

Ancestors may differ because of unpredictable factors such as mutation (Gould 2002). Alternatively, ancestral differences may reflect the different sequence of environments experienced during the divergence of the ancestral forms from *their* common ancestor (Price et al. 2000; Prunier et al. 2012). If the latter is important, a predictive theory of contingency should be possible to develop. This theory has two main components. First, we need to understand the reasons the ancestors differ, and second, we need to reconstruct how ancestors affect subsequent diversification. In this paper, we use this two-step approach to compare the visual pigments of two clades of birds that occupy a similar range of environments on different continents, while having drastically different plumage colors and diversity. Our goal is to ask if divergence in the visual system could contribute to the evolution of very different plumage colors in the two groups (Fig. 1).

**Fig. 1.**
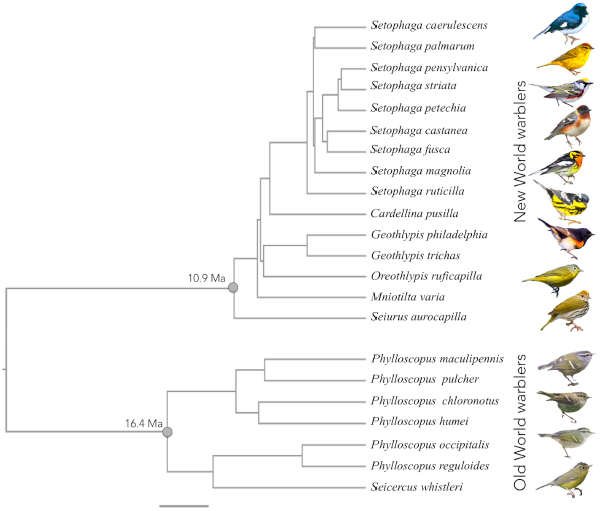
Time calibrated phylogeny of warbler species studied. Branch lengths are proportional to time and scale bar indicates 4 million years. We used a previously published phylogeny for New World warblers (Lovette et al. 2010). The Old World warbler phylogeny, connection between the two clades and absolute dates are from Price et al. (2014). The date of insert of the New World clade was estimated from an additional analysis following the methods of Price et al. (2014) that included *Seiurus aurocapilla* sequence. Mean node ages and corresponding 95% confidence intervals from a Bayesian analysis are as follows: New World warblers 10.89Ma [9.28-12.66]; Old World warblers 16.37Ma [14.30-19.51]; last common ancestor to New World and Old World warblers 29.66Ma [26.9-33.15]. Illustrations are examples of male individuals of a few species for each clade. From the top: *Setophaga caerulescens, S. palmarum, S. pensylvanica, S. castanea, S. fusca, S. magnolia, S. ruticilla, Oreothlypis ruficapilla, Seiurus aurocapilla, Phylloscopus maculipennis, P. humei, P. reguloides* and *Seicercus whistleri*.

We illustrate the general framework behind this study in Figure 2, where we consider parallelism, convergence and historical contingency as alternative evolutionary outcomes when ancestral forms come to occupy similar environments. Jablonski (in Pearce 2012) defines parallelism to be evolution of the same trait from the same ancestral form (Fig. 2a) and convergence the evolution of the same trait from different ancestral forms. However, unlike parallelism, convergence is often studied when an ancestral trait is retained in one lineage but lost and then regained in another (Fig. 2b). Parallelism and convergence are distinguished because parallelism is taken to emphasize that a limited set of genetic/developmental variants channel directions of evolution, whereas convergent evolution more strongly implicates a role for selection in directing evolutionary trajectories (Pearce 2012). Similar principles regarding the guiding roles of selection and development can be applied to contingency (Losos 2010). In this case, ancestral differences lead to different solutions in response to similar selection pressures, rather than converging to the same phenotype (Fig. 2c). As differences between ancestors accumulate, we expect parallelism to become increasingly less common compared to convergence (Conte et al. 2012; Fig. 2) and a greater role for contingency in limiting the extent of convergence or even promoting divergence (Gould 2002; Ord 2012).

**Fig. 2.**
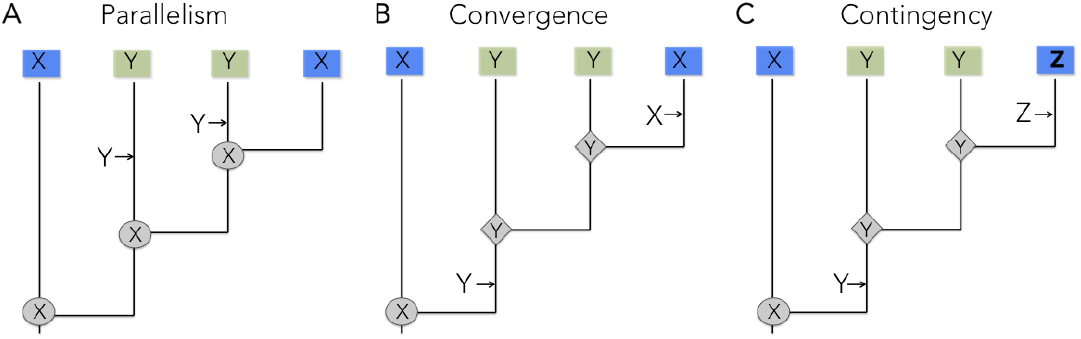
A framework for the study of parallelism (A), convergence (B), and contingency (C). Shading of squares indicates different environments, and X, Y, and Z are three different states of a trait in ancestors and their descendants. Arrows indicate transitions between states. In panel (C), the presence of trait Y in the ancestor results in further divergence when the ancestor re-enters the original environment, rather than convergence.

Discriminating between parallelism, convergence and contingency is difficult for two main reasons. The first is that similarity at the level of the trait may not reflect similarity in the underlying genetic mechanisms (Arendt and Reznick 2008; Pearce 2012). For example, different mutations may account for cases of apparent parallel evolution and conversely, the same mutation could account for cases of apparent convergent evolution (Arendt and Reznick 2008). Indeed, Rosenblum et al. (2014) define parallelism and convergence as the independent evolution of the same trait in different lineages with either the same (parallel) or different (convergent) molecular and developmental mechanisms, which is only testable once these mechanisms are understood. The second reason is that all interpretations rely on ancestral reconstructions. Ancestral reconstruction is especially challenging for phenotypic traits that show high evolutionary lability (Schluter et al. 1997), which is the case for many traits used in studies of convergent evolution. Gene sequences can often be reconstructed with a higher certainty than phenotypes. Thus, if the genotype can be linked to the phenotype, studying evolution at the genetic level can greatly improve assessments of parallelism, convergence and contingency. Opsin-based visual pigments provide one of the few systems where these approaches can be used (Chang 2003; Yokoyama 2008; Hunt et al. 2009).

Visual pigments consist of an opsin apoprotein bound to a light-sensitive chromophore. Birds possess four visual pigments mediating color vision, each encoded by the corresponding opsin gene: long-wavelength sensitive (LWS), medium-wavelength sensitive (RH2), and two short-wavelength sensitive visual pigments (SWS2 and SWS1). Within each pigment class, variation at key amino acid residues of the opsin protein causes differences in the visual pigment’s wavelength of peak absorbance (λ_max_), thereby affecting color perception (Bowmaker 2008; Hunt et al. 2009). In this system the relationship between genotype (sequence of opsin genes) and phenotype (λ_max_) can be ascertained by *in vitro* assays (Yokoyama 2008; Bowmaker 2008; Morrow and Chang 2010). We use the framework of Figure 2 to study the phenotypic evolution of the opsin genes in the Old World warblers (Phylloscopidae) and the New World warblers (Parulidae)

The Old World warblers and the New World warblers are small insectivorous leaf gleaning species breeding in the forests of Eurasia and North America, respectively (Price et al. 2000). The Old World warblers are sexually monomorphic and differ mostly in the number of unmelanized plumage patches present in the plumage, along with minor variation in carotenoid coloration (Price and Pavelka 1996). By contrast, the New World warblers are often sexually dichromatic and differ strikingly in carotenoid, structural and/or melanin pigments, with blues, reds and yellows all dominant features of the plumage of different species (Fig. 1). Based on a time-calibrated tree (Price et al. 2014) the two clades last shared a common ancestor in Asia around 30Ma. The New World warblers were derived from an ancestor that crossed the Bering Strait (Barker et al. 2004) about 15-20Ma (Price et al. 2014). Successive outgroups to the New World warblers are the New World sparrows, the buntings and the finches (Barker et al. 2004; Price et al. 2014), all of which are predominantly open-country omnivorous species. This suggests that as ancestors to the New World warblers diverged from their Old World counterparts, they passed though a finch-like, open-country species before moving back into forest habitats. This historical inference closely corresponds to the framework illustrated in Figure 2b and 2c, setting up the perfect system in which to test for historical contingency.

Why is it that New World warblers did not re-evolve dull plumages as this finch-like ancestor moved back into forest environments? While ancestral differences in features of the plumages themselves may contribute to the color differences between the New and Old World warblers (Price et al. 2000), here we focus on possible effects resulting from the evolution of the visual pigments. Many sexual selection models predict a role for visual perception in the evolution of colorful animal signals such as plumage. This idea is embodied in models for the evolution of mate choice, such as runaway sexual selection, the good genes and other models where a trait carries information on male quality, and in sensory drive models, where perception evolves in response to the environment (Andersson 1994; Boughman 2002; Horth 2007). As an important component of color perception, visual pigments could be at the basis of contingency in the evolution of animal signals such as plumage colors.

We surveyed the complete sequences of the four cone opsin genes across 22 species of New and Old World warblers. We focus on SWS2 evolution as it differs substantially both within and between the New World and Old World warblers while the other opsins are more conserved (Bloch 2014). Because evolution of SWS1 has been shown to be correlated with that of SWS2 (Hart and Hunt 2007), we study the evolution of this pigment as well. We investigated the evolution of the two short-wavelength sensitive visual pigments to address the following hypotheses:

1. *The null hypothesis*: Despite sequence evolution, SWS2 spectral tuning (λ_max_) has remained unchanged during the course of evolution of the New World warblers and Old World warblers from their common ancestor.
2. *Hypothesis of evolutionary convergence*: SWS2 λ_max_ is similar in the New World and Old World warblers, but different in ancestral forms, suggesting it has converged in response to environmental features of their forest habitats (Fig. 2b).
3. *Hypothesis of historical contingency:* SWS2 λ_max_ shifted as New World warblers diverged from Old World warblers passing through a finch-like ancestor that presumably occupied a different light environment and had different habits. Under this hypothesis evolution of SWS2 spectral tuning could affect color perception and contribute to plumage divergence in New World warblers (Fig. 2c).

## Materials and Methods

We sequenced the complete SWS2 opsin genes in both warbler clades and some outgroups, estimated ancestral states of SWS2 opsin gene sequences and finally, regenerated and measured the spectral sensitivity of warbler and ancestral SWS2 pigments. We combined our results with published measurements of SWS2 λ_max_ to test for an association of SWS2 spectral sensitivity with the light environment. We also studied the warbler’s SWS1 opsin genes and spectral sensitivity.

For the New World warblers, we consider species belonging to 6 out of the 14 genera (*Cardellina, Geothlypis, Mniotilta, Oreothlypis, Seiurus, Setophaga*) and follow previously established phylogenetic relationships for these species (Lovette et al. 2010). For Old World warblers, we include species belonging to the Old World-leaf warblers (family Phylloscopidae), a subset of the larger group commonly referred to as Old World warblers, which includes two genera, *Phylloscopus* and *Seicercus*. For this clade we follow phylogenetic relationships in Price et al. (2014).

### Tissue collection and opsin sequencing

For New World warblers we collected birds that died as a result of building collisions during migration in Chicago (Illinois, USA), and for Old World warblers we used RNA samples collected and processed by K. Marchetti (*pers. comm*.) in connection with other studies. We preserved eyes in RNAlater (Life Technologies) or liquid nitrogen in order to extract total RNA from the retinas of individual birds less than 2.5 hours post-mortem. Total RNA was extracted following TRIzol protocol (Life Technologies). In the 5 New World species with the highest RNA integrity (*Seiurus aurocapilla, Oreothlypis ruficapilla, Geothlypis philadelphia, Setophaga pensylvanica* and *S. palmarum*), we synthesized adaptor-ligated cDNA for use in RACE-PCR (rapid amplification of cDNA ends; SMART RACE system – BD Clontech). We used degenerate primers, designed based on available bird opsin sequences in Genbank, to amplify small coding sequence fragments. The resulting short fragments were used to develop 5’ and 3’ outward primers to use in RACE-PCR to obtain full-length opsin coding sequences. We then used the resulting full-length sequences for these 5 New World warbler species to design nested pairs of primers located in conserved regions of the 5’ and 3’ gene ends and/or UTRs of each opsin gene (see Table S1 for primer sequences). For all warblers, including the species used for the initial RACE-PCR, cDNA was synthesized from total RNA extracted from retinas for each individual using oligo-dT primers (with Qiagen’s Omniscript RT kit), and used to amplify the full coding sequences in conjunction with the nested primers we designed (Table S1). For all New World warblers and, when possible, Old World warblers, opsin sequences were amplified from more than one individual. Sequences are deposited in GenBank (accession numbers KM516225-KM516272).

### Ancestral reconstructions

We used available complete SWS2 opsin sequences from Genbank (SWS2 for canary, *Serinus canaria* - AJ277923, zebra finch, *Taeniopygia guttata* - AF222332, and chicken, *Gallus gallus* - NM205517) as well as our own outgroups (White-throated sparrow, *Zonotrichia albicollis*, Indigo bunting, *Passerina cyanea*, Yellow-bellied fantail, *Chelidorhynxhypoxantha*, and Goldcrest, *Regulus regulus* - KM516240/41/49/50) to reconstruct SWS2 sequence evolution. Phylogenetic relationships within the New World warblers were taken from Lovette et al. (2010), with all other relationships extracted from the tree of Price et al. (2014; see legend to Fig. 1). We performed parsimony reconstructions as implemented in Mesquite (Maddison and Maddison 2001)), as well Empirical Bayes (EB) ancestral reconstructions implemented in PAML (Nielsen and Yang 1998; Yang et al. 2000; Yang 2007). EB reconstructions can be based on nucleotide, amino acid and codon substitution models and use maximum likelihood estimates of branch lengths to assign the optimal character state at each amino acid site for all ancestral nodes. Because different types of models are sensitive to different assumptions, we performed ancestral reconstructions using nucleotide, amino acid and codon-based models and, where applicable, used likelihood ratio tests to choose the best fitting models for each type (Chang et al. 2002). We compared the sequences reconstructed under each model to check for the robustness of the ancestral states of all nodes and used posterior probabilities to determine the most likely protein sequence at each node (Chang et al. 2002).

### In vitro regeneration of visual pigments and spectral analyses

From the opsin sequence data, we identified all variants with at least one non-synonymous substitution as candidates for shifts in λ_max_. We did not express the SWS2 pigment of *S. whistleri*, which had a single valine to isoleucine substitution at site 166, as this substitution is not likely to cause a significant change to the physiochemical properties of the visual pigment (Shyue et al. 1998). The complete coding sequences of selected opsins were cloned into the p1D4-hrGFP II expression vector (Morrow and Chang 2010). These constructs were used to transiently tranfect cultured HEK293T cells using Lipofectamine 2000 (Invitrogen; 8 □g of DNA per 10-cm plate). Cells were harvested 48 h post-transfection and opsins were regenerated through incubation in 5 μM 11-cis-retinal generously provided by Dr. Rosalie Crouch (Medical University of South Carolina). All visual pigments were solubilized in 1% N-dodecyl-D-maltoside (DM) and immunoaffinity purified using the 1D4 monoclonal antibody (Molday and MacKenzie 1983), as previously described (Morrow and Chang 2010; Morrow et al. 2011). We used glycerol buffers to improve the stability of short wavelength-sensitive visual pigments during expression (Starace and Knox 1998). Purified visual pigment samples were eluted in 50mM sodium phosphate buffer (0.23% NaH_2_PO_4_, 0.43% Na_2_HPO_4_, 0.1% DM, pH 7).

The ultraviolet-visible absorption spectra of all purified SWS2 visual pigments were recorded at 25°C using a Cary4000 double-beam absorbance spectrophotometer (Varian) and quartz absorption cuvettes (Helma). All λ_max_ values were calculated after fitting data from multiple absorbance spectra of each visual pigment to a standard template for A1 visual pigments. This involved a baseline correction of raw absorbance spectroscopy data, then matching the slope of the long-wavelength arm to an appropriate Govardovskii template, as described elsewhere (Govardovskii et al. 2000). This process allowed for a more accurate estimation of a λ_max_ value for each visual pigment in order to facilitate the identification of any spectral shifts between variants. The λ_max_ values we present correspond to the average of three separate measurements of the absorbance spectrum for each pigment expressed. SWS2 visual pigments were bleached for 60 seconds using a Fiber-Lite MI-152 150-Watt Fiber Optic Illuminator (Dolan-Jenner), causing their λ_max_ to shift to ~380 nm, characteristic of the biologically active meta II intermediate. We calculated difference spectra by subtracting these light-bleached spectra from respective dark spectra. Since the λ_max_ of SWS1 pigments is similar to that of the light-bleached intermediate, an acid bleach (HCl to 100mM final concentration) was performed, causing a shift to ~440nm instead.

In order to assess reliability, some pigments were expressed a second time (see Table 1, 5^th^ column). Standard errors between replicates were comparable to within replicate measurements.

**Table 1.**
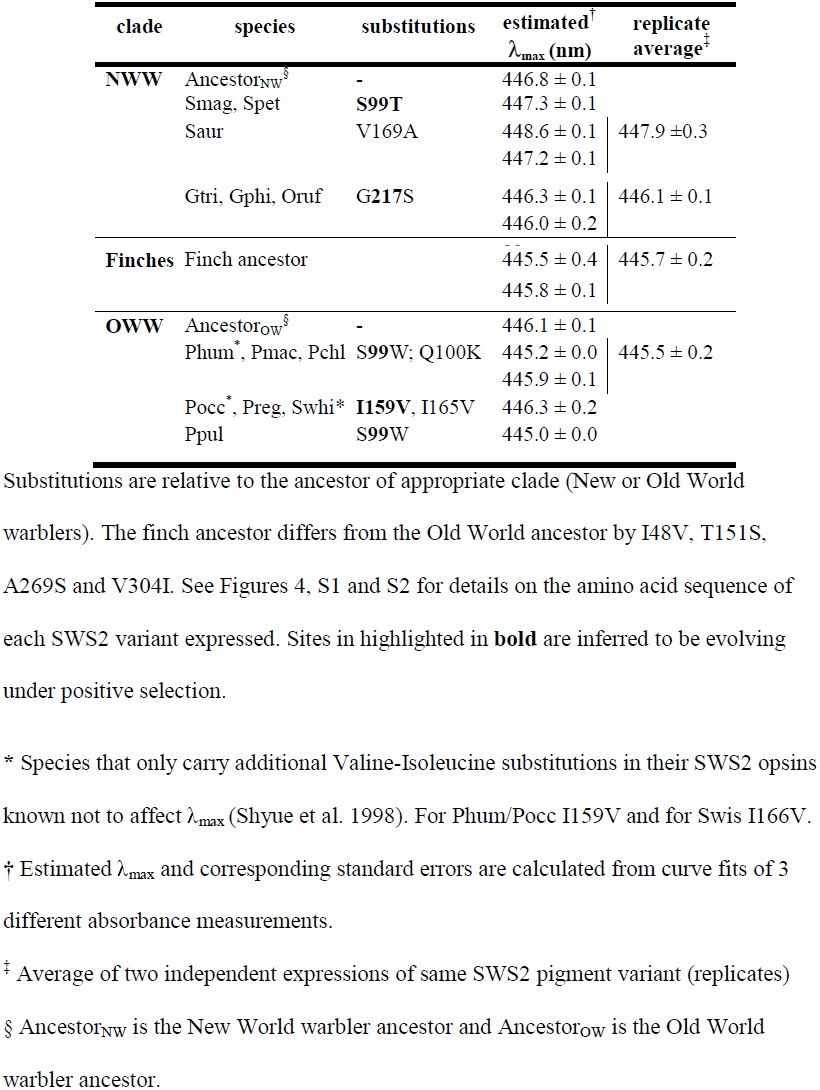
Spectral sensitivities for SWS2 visual pigments expressed *in vitro*

### Site-directed mutagenesis

The inferred ancestral sequence to the Old World warblers and the finches is not present in any of the extant species. We thus synthesized SWS2 pigments of these ancestors using site-directed-mutagenesis following QuikChange (Stratagene) protocols and using *PrimerX* software (http://www.bioinformatics.org/primerx/index.htm) to design mutagenesis primers. To recreate the Old World warbler ancestor’s sequence we used *Phylloscopus reguloides* as template, as it only has one substitution relative to the inferred ancestral sequence. We designed 100% complementary mutagenesis primers to introduce a V159I mutation into *P. reguloides* (forward primer CTGGGCTGCGCCATCACCTGGATCTTC, reverse primer GAAGATCCAGGTGATGGCGCAGCCCAGC). In the same way, to synthesize the finch ancestor’s SWS2 we designed primers to introduce an L49V mutation into *Geothlypis trichas* (forward primer GTTCCTGCTGGTGGTGCTGGGCGTGC, reverse primer GCACGCCCAGCACCACCAGCAGGAAC). Mutagenesis was performed in the TOPO vector (Invitrogen) following cycling conditions provided in the QuikChange protocol (Stratagene). To express these ancestral pigments *in vitro* and measure their spectral sensitivity we proceeded as described above for the naturally occurring SWS2 variants.

### Visual pigment molecular evolution

We used codon-based site models as implemented in PAML to identify sites evolving under positive selection (Yang and Bielawski 2000; Yang 2007). Here estimates of Ω = *d*_*N*_/*d*_*S*_ for each site are calculated in a maximum likelihood framework under various models that allow for different levels of heterogeneity in ω. The M0 or “one-ratio” model is the simplest model, assuming the same ω for all sites in all branches of the provided tree. Two nested pairs of models evaluate evidence for sites evolving under positive selection. In each pair the parameter rich model, that allows for an additional category of sites with ω > 1, is compared to a simpler model that does not. A likelihood-ratio test (LRT) is used to evaluate whether the more complex models (M2 and M8) fit the data better than the simpler models (M1 and M7 respectively). To minimize the possibility of reaching local optima we ran all models with different starting ω values (ω=0.0, 0.5, 1.0, 1.5, 2.0, 5.0). Finally, when models accounting for positive selection fit the data significantly better by the LRT, we used a Bayes empirical Bayes analysis, also implemented in PAML, to identify sites evolving under positive selection (Yang and Bielawski 2000; Yang 2005).

### Correlated evolution: SWS2 λ_max_ association with habitat

To test for an association between SWS2 spectral tuning and habitat we combined our data for New World and Old World warblers with all species whose SWS2 λ_max_ has been measured so far. This added 12 passerine species, 5 of which are finches, and 11 non-passerines (Table S2). Among these additional species, all but the zebra finch (Yokoyama et al. 2000) have been studied using microspectrophotometry (MSP) on retinas (Bowmaker et al. 1997), which does not require opsin genes to be sequenced, but λ_max_ estimates are less precise than the *in vitro* expression we use here. We classified habitat into three easily quantified categories, as these have been shown to follow an important axis of light quality variation in forests (Endler 1993; sources are in Table S2). We treated SWS2 λ_max_ as a continuous dependent variable and assessed associations with habitat scored on a 3 point scale: 1) “Forest understory” for species that spend significant portion of time foraging on the ground or undergrowth and 2) “Arboreal” for species foraging in the forest higher than 1m (Lovette and Hochachka 2006). Following Lovette and Hochachka (2006) we did not separate species into low-mid-or high canopy foragers, because these partitions are ill defined. 3) “Open” for non-forest species that forage out in the open, in swamps or wetlands. Birds that recently adapted to urban life as human commensals (e.g. *Turdus merula*) were classified according to their native/ancestral habitats. Of the 15 species of New World warblers we studied, one is classified as occupying the forest ground/undergrowth (the ovenbird, *Seiurus aurocapilla*), 10 are arboreal, and four fall in the open habitat category (Table S2). We classified all the Old World warblers we studied as arboreal (Price et al. 2000; Ghosh-Harihar and Price 2014).

We conducted ordinary and phylogenetic least squares regression of log(λ_max_) values against foraging habitat (Orme et al. 2013). For phylogenetic correction, we used the phylogeny of Jetz et al. (2012, maximum clade credibility of the first 1,000 trees and the "Hackett backbone" downloaded from birdtree.org, except we replaced the New World warbler clade in that tree with the one from Lovette et al. (2010). We used the (Jetz et al. 2012) tree because it contains all species for which spectral tuning has been measured (the Old World warbler topology is identical, and branch-lengths very similar to the one in Price et al. (2014)). Phylogenetic least squares regression is identical to the commonly used independent contrasts model, except that it allows for an adjustment in branch lengths to optimize the Brownian motion assumption. Because the independent variable (habitat) is categorical and not numeric, we assessed significance of associations using ANOVA. We constructed the ANOVA model by adding two dummy columns as independent variables, which contrasted the (1) first two categories against the third, and (2) the first category against the other two. We then subjected the P value to the ordered ANOVA test (Rice and Gaines 1994).

## RESULTS

### Molecular evolution

A total of 11 amino acid sites vary in the SWS2 genes of New World and Old World warblers (Figs. 3, S1). Three amino acid sites vary within the New World warblers and four within Old World warblers, with several cases of parallel evolution and reversals (Figs. 3, 4, S2). With respect to parallel evolution, substitution S99T (using bovine rhodopsin residue numbering) is inferred to have happened twice in the tips of the New World warbler tree, in the lineages leading to *Setophaga magnolia* and *S. petechia*, and substitution I159V is found twice in the Old World warblers. Reversals are present at site 100 in Old World warblers (Q100K back to K100Q in *P. pulcher*) and site 217, which reversed from glycine in the ancestor of sparrows back to serine in the branch leading to the *Geothlypis/Oreothlypis* clade in the New World warblers (Fig. S2). Using maximum likelihood site-models in PAML, we identified two sites as having evolved under positive selection within New World warblers and two within the Old World warblers (Fig. 3, Tables 1, S3). Site 99, which underwent different substitutions in each clade, is inferred to have evolved under positive selection in both clades (Fig. 3, Tables 1, S3).

**Fig. 3.**
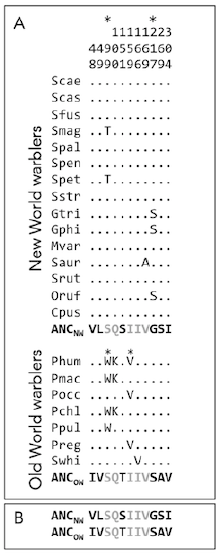
Alignment showing only variable amino acid residues of the SWS2 coding sequences for the New World warblers and the Old World warblers based on full coding sequences. Numbers correspond to amino acid positions standardized by the bovine rhodopsin (GenBank M21606). Refer to Figure S1 for the precise location of each substitution relative to transmembrane domains. Species names are abbreviated as the first letter of the genus and the three first letters of the species (i.e. *Setophaga castanea* is Scas, see Fig. 1 for full species names). All variable sites are shown relative to the inferred ancestor of each clade: ANC_NW_ for New World and ANC_OW_ for Old World warblers, as obtained by likelihood/Bayesian methods. Dots indicate the identity of the amino acids with the ancestor sequences at each site, thus species that only have dots match the ancestral amino acid sequence at all sites. As highlighted in panel (b), the SWS2 sequence of both ancestors differs between clades. Positions in grey correspond to residues that have the same amino acids in the ancestors of New World and Old World warblers and residues in black are those that differ between both ancestors and thus with fixed differences between both clades. *Sites identified as evolving under positive selection within clades (see Table S3).

**Fig. 4.**
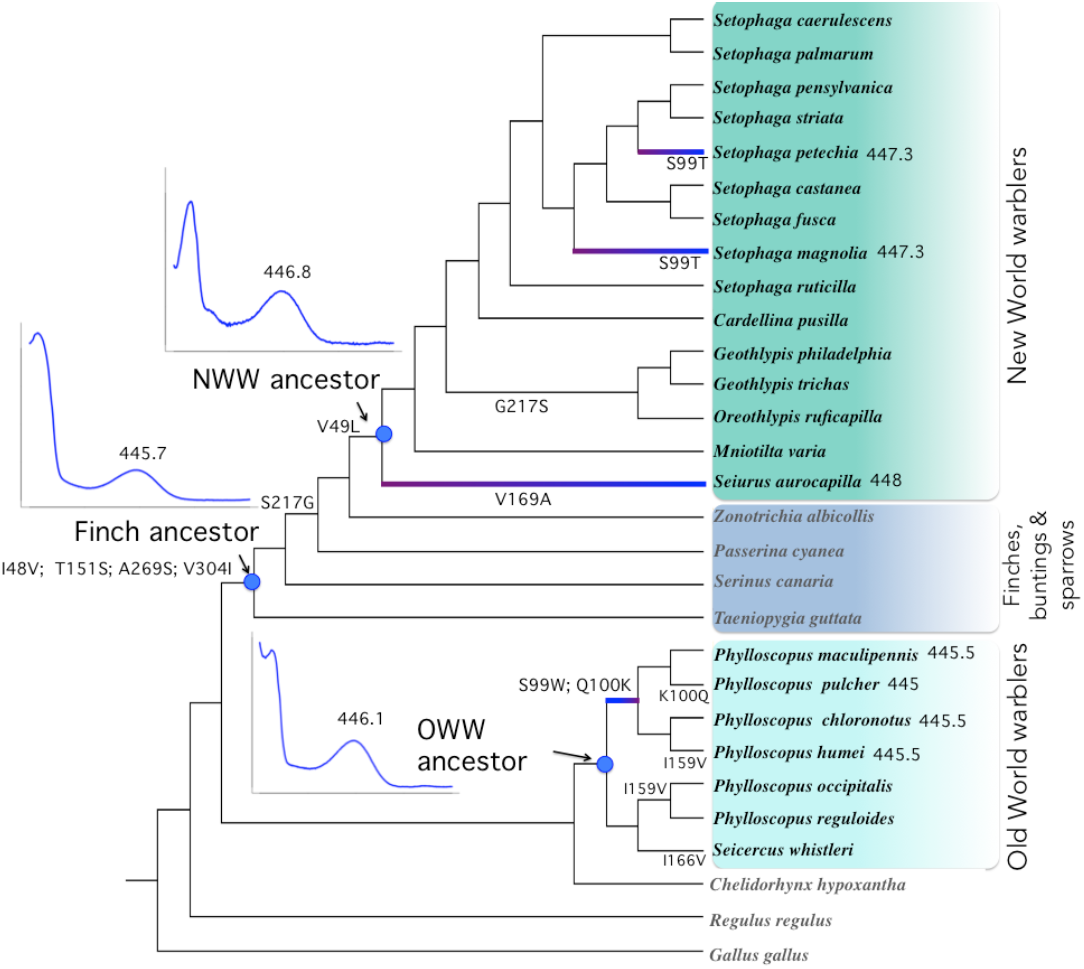
Cladogram of SWS2 sequence evolution in New World warbler and Old World warbler species, with the topology from Price et al. (2014) and Lovette et al. (2010), as in Figure 1. Indents show SWS2 absorbance spectra, and their corresponding λ_max_ value, given the inferred sequences for the ancestors of New World and Old World warblers, as well as the finch ancestor (axis scales for these graphs are the same as Fig. 5). Highlighted branches illustrate spectral shifts associated with warbler evolution, as listed in Table 1, as well as their direction. SWS2 λ_max_ values for these branches are shown next to species names. Substitutions shown for each edge correspond to the states with the highest posterior probabilities from likelihood/Bayesian ancestral reconstructions. The deepest node in this tree has the following inferred amino acid composition at the relevant sites: I48, V49, S99, Q100, T151, I159, I166, V169, S217, A269, V304. Posterior probabilities associated with the ancestral reconstruction of amino acid sequences are shown in Fig. S2.

All ancestral reconstruction models were mostly in agreement except for one minor difference identified below. Model comparisons to determine the best fitting nucleotide, codon and amino acid models are summarized in Table S4; in Figure 4 we show the sequence in which substitutions accumulated at all nodes. We infer the last common ancestor to the New World and Old World warblers to have had the same amino acid sequence as the Old World warbler ancestor (all posterior probabilities > 0.94 except for S269 = 0.775; see Fig. S2). In fact, the chicken (a distant non-passerine) has the same amino acids at these 6 sites as the Old World warbler ancestor. This implies that the 6 SWS2 amino acid differences between the New World and Old World warblers were all substituted along the lineage leading to the New World warblers. Four substitutions (I48V, T151S, A269S, V304I) accumulated early in the history of divergence, before finches (Fringillidae and Estrildidae) split from the New World warblers. Two substitutions (S217G, V49L) occurred along the branch from the finch ancestor to the New World warbler ancestor. There is a minor disagreement among reconstruction models for substitution S217G. Codon and nucleotide models concurred that this substitution happened on the branch shown on Fig. 4 and Fig. S2, but amino acid based models inferred it occurred along the same branch as V49L.

### Spectral tuning

We found that, despite having 6 inferred amino acid differences, the ancestors of the two warbler clades have similar λ_max_ values differing by 0.7nm in their point estimates (Figs. 4, 5 and Table 1). Furthermore, we found only small spectral tuning differences within clades: point estimates vary by up to 2nm within the New World warblers and 1.3nm within the Old World warblers (Table 1).

**Fig. 5.**
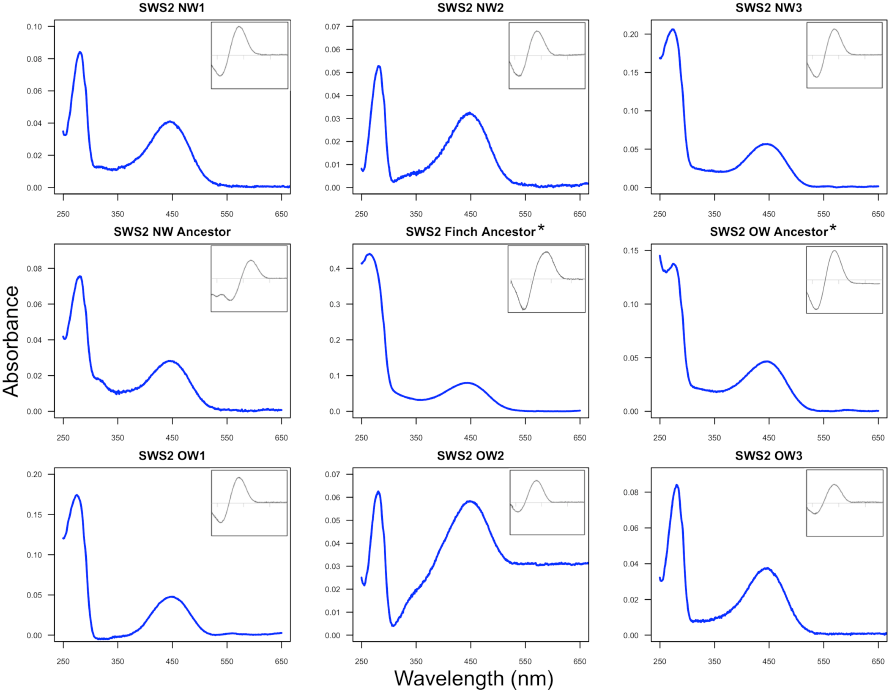
Absorbance spectra of the regenerated warbler SWS2 pigments expressed *in vitro*. Main graphs show dark spectra and insets correspond to dark-light difference spectra. The x-axis for insets has the same range as main graph in all cases. Variant name correspondence: NW1 (Smag, Spet, λ_max_ 447.3nm), NW2 (Saur, λ_max_ 448nm), NW3 (Gtri, Gphi, Oruf, λ_max_ 446.1nm), OW1 (Phum, Pmac, Pchl, λ_max_ 445.5nm), OW2 (Pocc, Preg, Swhi, λ_max_ 446.3nm) and OW3 (Ppul, Xmax 445nm). See Table 1 for further details. *Ancestors recreated by site-directed mutagenesis. New World warbler ancestor λ_max_ 446.8nm, Old World warbler ancestor λ_max_ 446.1nm, and finch ancestor λ_max_ 445.7nm. Note that the results are not normalized so the y-axes of different graphs have different scales due to absolute differences in expression.

The SWS2 λ_max_ values within the New World and Old World warbler clades are very similar, with, for example, *Geothlypis trichas* (New World) and *Phylloscopus occipitalis* (Old World) having near identical point estimates (Table 1), despite their sequence differences. Such similarity may reflect either inheritance through the common ancestor or convergence following a shift in spectral tuning in the finch ancestor. To test for these alternatives, we used site-directed mutagenesis to reconstruct the finch ancestor’s SWS2 and found it has a λ_max_ = 445.7nm ± 0.2 (Table 1), extremely similar to the warbler ancestors. The short-wavelength shifts in finches evidently accumulated after they diverged from the warbler lineage, and the similarity within the warbler clades appears to be a consequence of shared ancestry, not convergence.

### SWS1 evolution

We showed above that SWS2 λ_max_ is very similar among warblers. Hart and Hunt (2007) found that across all birds variation in SWS2 λ_max_ can be partially explained by the λ_max_ of the SWS1 pigment, with SWS2 λ_max_ tuned to shorter wavelengths in species with a UV-shifted SWS1. Most passerines, including all those considered in Hart and Hunt (2007), have a UV sensitive SWS1, with exceptions in basal clades (Coyle et al. 2012; Ödeen et al. 2012). The New World warbler inferred ancestor SWS1 sequence was maintained in most extant warblers. *Seiurus aurocapilla* is the only species with non valine-isoleucine SWS1 substitutions (M109L and E280D). When expressed *in vitro*, we found the New World warbler ancestral SWS1 and *S. aurocapilla’s* SWS1 had similar λ_max_ values (365.1 ± 0.1 nm and 364.8 ± 0.2 nm, respectively; Fig. 6a & b). In a similar way, we found SWS1 is very conserved across Old World warblers with a λ_max_ of 362.6 ± 0.3 nm (Fig. 6c), indicating all warblers have UV sensitive SWS1 pigments, and experience minimal variation in λ_max_. We computed the correlation between SWS1 λ_max_ and SWS2 λ_max_ using only the Passerines in Hart and Hunt (2007) and Coyle et al. (2012) and found no association (Pearson *r*_*s*_ = 0.36, P = 0.28, N = 11; data in Table S2). This implies that SWS2 spectral tuning in Passerines varies beyond any co-evolutionary process with SWS1.

**Fig. 6.**
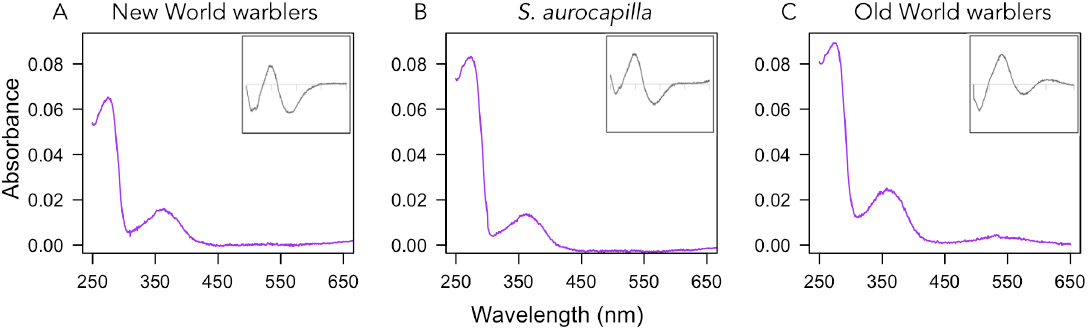
Absorbance spectra of the SWS1 pigments regenerated *in vitro*. (A) SWS1 shared by all New World warblers except *S. aurocapilla* (365.1 ± 0.1 nm). (B) SWS1 of *S. aurocapilla* (364.8 ± 0.2 nm), which carries substitutions M109L and E280D relative to other warblers in its clade. (C) SWS1 pigment of Old World warblers (362.6 ± 0.3 nm). Main graphs show dark spectra and insets correspond to dark-bleached difference spectra (after acid bleaching). *x*-axis for insets has the same range as main graph in all cases.

### Adaptive significance of SWS2 phenotypic variation

Across passerine species, SWS2 λ_max_ is correlated with foraging habitat (Figure 7; in an ordered ANOVA F_2,31_ = 17.99, P <0.0001; phylogenetic control using phylogenetic least squares regression P =0.0016; see Fig. S4). Species that forage in the forest understory have relatively long-wavelength shifted SWS2 visual pigments compared to those in more open habitats, matching the spectral properties of the available light in these habitats (Fig. 7B). We added non-passerine species for which data is available (Hart and Hunt 2007) in a model that included SWS1 λ_max_ as a covariate (Table S2). The association between SWS2 spectral tuning and the environment still holds on this larger phylogenetic scale (phylogenetically corrected analysis, Pλ0.0001, with a significant effect of SWS1 λ_max_ P=0.016; non-passerines alone P= 0.001, SWS1 λ_max_ P=0.013, for details see Fig. S6 and Table S2). It is also worth highlighting that despite the small differences in spectral tuning among New World warblers (2nm at the most), the one species that inhabits the forest understory in our dataset (*Seiurus aurocapilla*) has the most long wavelength shifted SWS2 λ_max_, in accord with the general association across all species. Since most of the data we used from previous studies were obtained using microspectrophotometry (MSP) we made sure that the contrast between techniques was not driving the relationship we found. Using only MSP data, the relationship between foraging habitat and SWS2 spectral tuning is still significant (Table S2; N = 11 species, P = 0.0002 based on an ordered ANOVA; phylogenetic control, P = 0.001).

**Fig. 7.**
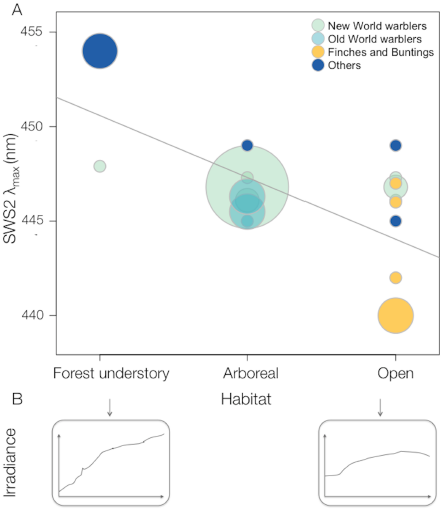
Scatter plot of SWS2 λ_max_ against habitat category in Passerine birds. Circle size corresponds to number of species and circle shade indicates the clade as detailed in the figure legend. Breeding season foraging habitat categories correspond to (1) species foraging in the forest understory, on or close to the ground, (2) all the remaining strata of the forest and (3) outside forests in the open, based on 22 warbler species from this study and 12 additional species of passerines (Table S2). The regression line, y= 450.6-3.3x, corresponds to a simple regression where habitats are given numerical values (0, 1, 2). Significance was calculated based on an ordered ANOVA where the values are considered categorical (P <0.0001, phylogenetic correction P <0.0016; details as Supplementary information). Bottom insets are irradiance spectra from 400-700nm for “small gaps” and “large gaps” respectively, taken from (Endler 1993). A cladogram illustrating the correspondence between SWS2 spectral tuning and foraging habitat is in Fig. S4.

## DISCUSSION

The New World warblers were derived from the Old World ancestor, apparently passing through a finch or bunting-like ancestor in the process (Fig. 4). Many finches and buntings are colorful and dimorphic (Stoddard and Prum 2008). One possible reason is that open country habitats favor visual cues (Crook 1964). We considered that this should not only affect the evolution of colors directly, but also that the visual system would diverge as a consequence of passing through a finch-like ancestor. This is supported by a correlation between habitat and SWS2 tuning among present-day species (Fig. 7, S4). When the New World warbler ancestor re-entered forest habitats and an insectivorous niche, some ancestral features appear to have been retained during diversification. For example, several New World warblers eat fruit, whereas none of the Old World warblers do (Price et al. 2000). Here, we asked if SWS2 tuning evolved during passage through the inferred open country ancestor, and if this then left a lasting impression on the New World warblers.

Altered SWS2 spectral tuning in the New World warblers seemed especially likely considering that SWS2 accumulated 6 amino acid substitutions along branches leading to the New World warblers (Fig. 4). Further, two of the 6 sites (residues 49 and 269) have been studied using site-directed mutagenesis and shown to cause spectral shifts in SWS2 in the green anole and goldfish (Yokoyama 2003), as well as LWS in human (Asenjo et al. 1994) and bovine rhodopsin (Chan et al. 1992). However, we found that SWS2 λ_max_ is similar in the New World and Old World warblers as well as the reconstructed ancestors to these clades. Combined with the general similarity across clades in the tuning of the other opsins (Bloch 2014) this appears to rule out spectral tuning differences as a contributing factor to the differences in plumage coloration of the two groups.

Similarity across clades in SWS2 spectral tuning could be a result of shared ancestry or evolutionary convergence. With respect to convergence, progression to a finch-like form during the colonization of the New World, accompanied by the inferred four substitutions in the SWS2 opsin could have resulted in a spectral shift towards shorter wavelengths, followed by two additional substitutions that could have shifted SWS2 λ_max_ back to the longer wavelengths characteristic of the warblers. However, our reconstruction of the ancestral opsins implies that this was not the case and λ_max_ similarity between the clades apparently results from inheritance through a common ancestor, not convergence. Two major questions arise out of these results. First, given present day finches have short-wavelength shifted SWS2 (Fig. 7), why was spectral tuning not shifted in the inferred finch-like ancestral form SWS2? Second, if not tuning, what are the selective forces responsible for the substitutions present in the New World warbler lineage? We conclude by considering the implications of our results for the study of historical contingency and convergence more generally.

### What is driving SWS2 spectral tuning evolution?

Previously demonstrated correlates of spectral tuning in the terrestrial environment have been related to the colors of frequently encountered objects. They generally reflect single case studies and include both detection of prey (Regan et al. 2001) and conspecifics (Arikawa et al. 2005; Briscoe et al. 2010). In birds, the only correlate has been that of SWS1 λ_max_ (as inferred from DNA sequences rather than directly measured) with UV plumage reflectance in the fairy-wrens (Ödeen et al. 2012). However, in the aquatic environment visual pigment differences across species have been related to gradients of light intensity (Lythgoe 1984; Partridge and Cummings 1999; Seehausen et al. 2008). Variation matches not only light intensity, but can also be related to differences in the spectral composition of light (Partridge and Cummings 1999; Seehausen et al. 2008).

Correlates with terrestrial light environments have been much more difficult to demonstrate than in aquatic environments (Lythgoe 1979). Here we found that across all passerines, there is a highly significant association between SWS2 λ_max_ and the inferred light environment (Fig. 7). Species in environments with less short-wavelength light have relatively long-wavelength shifted SWS2 λ_max_. This relationship follows changes in the spectrum of light in these habitats: in the lower strata of the forest, light is relatively rich in mid to long wavelengths, because short wavelengths (“blue” and “UV”) are filtered out as light passes through trees, and in contrast, open habitats are relatively richer in short-wavelength light (Endler 1993; Gomez and Thery 2007). The simplest explanation is that SWS2 λ_max_ spectral tuning improves signal/noise ratio, which enables better contrast detection (Lythgoe 1979).

One caveat to this result is the difference in the methods used to measure SWS2 λ_max_ between previous studies and our own. Except for the Zebra finch, all the SWS2 spectral tuning information apart from our study was measured by microspectrophotometry (MSP), which is characterized by larger error than *in vitro* measurements of heterologously expressed visual pigment λ_max_ (Table S2). However, for most species and particularly the ones with extreme SWS2 λ_max_ values, differences are beyond the errors reported by the original studies (Table S2). An additional factor that needs to be considered is the presence of oil droplets, which could affect photoreceptor sensitivity. Oil droplets are found in the inner segments of cones, in the path of the light before it hits the visual pigment containing outer segments (Goldsmith et al. 1984). These droplets contain carotenoid pigments that filter short-wavelength light and thus, act as long-pass cutoff filters that narrow the spectral sensitivity and can shift the λ_max_ of the photoreceptor that contains them (Goldsmith et al. 1984). Little information is available on how oil droplet absorbance varies across species. We know that oil droplet pigment content, and thus spectral properties, change across the retina (Knott et al. 2010), and are sensitive to ambient light conditions (Hart et al. 2006) and carotenoid content in the diet (Bowmaker et al. 1993; Knott et al. 2010). Existing studies suggest the spectral properties of the oil droplets associated with SWS2 cones – “C-type”-are similar across passerines, and the range of variation within species approximates differences between species (Begin and Handford 1987; Bowmaker et al. 1993; 1997; Hart et al. 2000; 2006). It is even possible that SWS2 droplets do not contain enough filtering pigment to act as true cut-off filters (Bowmaker et al. 1993). Independent of the properties of oil droplets, the association between SWS2 spectral tuning and habitat suggests environmental pressures are shaping the evolution of this visual pigment.

The short-wavelength shifted SWS2 pigments in modern finches raises the question of why SWS2 λ_max_ was not short-wavelength shifted in the inferred finch-like ancestor. One reason may be that the habitat occupied by ancestral birds always favored an SWS2 λ_max_ around 446 nm. A second possibility is that opsin sequence evolution is slow in response to changing light environments, requiring long waiting times for the appropriate mutations to arise and be fixed. The probability of fixation of a new mutation is very low when selection is weak (Haldane 1927), implying many new mutations at the same site are required before one becomes established. In this case, little evolutionary change may have occurred in the finch transitional form before selection pressures again favored the warbler phenotype over that of the finch. Moreover, the visual system shows a great deal of plasticity, including neurological and physiological mechanisms, such as chromatic adaptation, which leads to color constancy under different illuminants (Foster 2011), and plasticity in the above-mentioned oil droplets (Bowmaker et al. 1993; Knott et al. 2010). Such plasticity is likely to lower selection coefficients on new mutations, especially if shifts in λ_max_ are small.

### Why have substitutions accumulated in SWS2?

The SWS2 substitutions are likely to have been fixed for adaptive reasons. First, according to molecular tests using PAML, at least some of the sites we have detected have been subject to positive selection. Second, no substitutions at these positions occurred throughout the long history up to the ancestor of the Old World warblers from the non-passerine split; if the substitutions had no effect on phenotype this period of long stasis is very unlikely. Third, parallel amino acid substitutions and reversals are present within the warbler clades (Fig. 4).

Assuming the fixed substitutions between the clades are adaptive, selection pressures could relate to the small differences in tuning we found or to other opsin functions. Theoretically, even small spectral shifts can affect color discrimination when they co-occur with changes in pigment density in the cones (He and Shevell 1995), an entirely unexplored aspect of avian vision. Small differences in visual pigment sensitivity are known to affect perception in humans (Mollon 1992). Variation of 3-5 nm caused by a polymorphism at position 180 of human LWS has a significant impact on color discrimination (Mollon 1992), causing differences large enough to lead subjects to score differently in standardized color matching tests (Sanocki et al. 1994). Whether the small shifts we found among warblers affect color vision is not known.

Alternatively, selection pressures may relate to other aspects of opsin function. For example, a study of rhodopsin in the echidna, *Tachyglossus aculeatus*, highlighted a series of amino acid substitutions that altered several aspects of visual pigment function, including the rate of retinal release and hydroxylamine sensitivity, sometimes without significant changes to λ_max_ (Bickelmann et al. 2012). These are aspects of visual pigment function that could also help organisms adapt to their light environments (Sugawara et al. 2010).

Eleven sites in SWS2 sequences have been altered during warbler evolution. While sites 49 and 269 have previously been implicated in the spectral tuning of RH2, SWS1 and SWS2 (Yokoyama 2008); (Takenaka and Yokoyama 2007) as well as LWS (Chan et al. 1992), the 9 remaining sites have either never been studied in isolation, or have not been implicated in spectral tuning in either SWS2 opsins (Cowing et al. 2002; Takahashi and Ebrey 2003; Yokoyama 2003; Yokoyama et al. 2007), or any other opsin class (Yokoyama 2000; Hunt et al. 2009). It is possible that substitutions at these sites have consequences for other aspects of opsin function. For example, sites 99 and 100 are part of transmembrane helix 2 (TM2; Fig. S1), which contains residues involved in establishing the hydrogen bonding network in the chromophore binding pocket (Palczewski et al. 2000). Sites 151, 159, 165 and 169 are part of TM4 (Fig. S1), which contains the hypothesized dimerization surface of rhodopsin (Fotiadis et al. 2003; Liang et al. 2003). Finally, site 217 is situated in the region of TM5 near a possible retinal exit site following activation (Hildebrand et al. 2009), where substitutions can alter the rate of retinal release (Piechnick et al. 2012; Morrow and Chang *in prep*).

### Contingency and convergence

Assessment of parallelism, convergence and contingency requires ancestral reconstructions (Fig. 2). Based on reconstruction of the finch ancestor, and assuming other steps we have not reconstructed did not involve spectral shifts, the similarity in spectral tuning in the Old World and New World warblers is a result of inheritance through a common ancestor, and not convergence. One implication from this study, deserving further investigation, is that finches and buntings appear to have evolved short wavelength-shifted spectral tuning in parallel (Figs. 4, S4).

Despite much research into the genetic basis of convergence over the last decade, studies that actually demonstrate the ancestor was different than the derived form are rare (e.eg. Manceau et al. 2010; Liu et al. 2010). Recent striking examples of repeated sequence evolution of the genome across distantly related groups in similar environments are strong circumstantial evidence for convergence (Castoe et al. 2009; Parker et al. 2013), but these studies alone cannot rule out parallelism, or even contingency, which requires estimates of the phenotype in ancestors, as well as present day species. Some studies have partially circumvented this difficulty by inferring function from sequence changes (Sugawara et al. 2005; Hofmann et al. 2012). Here, we were able to perform a complete test for convergence and contingency, because in this system we can link opsin protein sequence directly to spectral tuning.

In summary, we have compared SWS2 visual pigment tuning among and within two clades of birds in order to assess a possible role for color perception in color diversification. We found that spectral tuning has remained similar through the divergence of New World and Old World warblers even as opsin sequences have evolved, suggesting other features of the opsins besides tuning have driven their sequence evolution. Those features remain to be determined. When they are, it should be possible to assess whether passage through a finch-like form resulted in divergence of the visual system, with potential consequences for divergence in color. More generally, the two-step assessment of contingency involves asking why ancestors differ and then how those differences contribute to lasting differences between clades. Such methods should be more generally applicable across a wide range of phenotypes, as we learn more about the history of groups, and are able to relate genotype to phenotype.

## ACKNOWLEDGEMENTS

We especially thank I. van Hazel for her invaluable help with visual pigment *in vitro* expression. We thank K. Marchetti for assistance and samples, J. Endler and D. Jablonski for valuable discussion and comments, as well as the Associate Editor and two anonymous reviewers for a careful and constructive review of this manuscript. We gratefully acknowledge the Field Museum of Natural History and the Chicago Bird Collision Monitors for all their help collecting warbler specimens. This work was supported by the National Institute of Health NRSA 1F31EY020105 (to NIB) and National Science Foundation DEB 1209876 (to NIB), and a Natural Sciences and Engineering Research Council Discovery Grant (to BSWC).

